# COMMIT: Consideration of metabolite leakage and community composition improves microbial community models

**DOI:** 10.1101/2021.06.02.446851

**Authors:** Philipp Wendering, Zoran Nikoloski

## Abstract

Composition and functions of microbial communities affect important traits in diverse hosts, from crops to humans. Yet, mechanistic understanding of how metabolism of individual microbes is affected by the community composition and metabolite leakage is lacking. Here, we first show that the consensus of automatically generated metabolic models improves the quality of the draft models, measured by the genomic evidence for considered enzymatic reactions. We then devise an approach for gap filling, termed COMMIT, that considers exchangeable metabolites based on their permeability and the composition of the community. By applying COMMIT with two soil communities from the *Arabidopsis thaliana* culture collection, we could significantly reduce the gap-filling solution in comparison to filling gaps in individual models. Inspection of the metabolic interactions in the soil communities allows us to identify microbes with community roles of helpers and beneficiaries. Therefore, COMMIT offers a versatile automated solution for large-scale modelling of microbial communities for diverse biotechnological applications.

## Introduction

Microbial communities have been extensively studied due to their importance in ecology (1, 2), human health (3, 4), and biotechnological applications (5). It has also been suggested that microorganisms have particular roles in a community; for instance, the Black Queen (BQ) hypothesis (6) suggests the existence of BQ functions, such as production of membrane-permeable products, that are essential for members termed helpers, but unavoidably available to other community members, termed beneficiaries. These roles are achieved by active transport and/or leakage of diverse metabolites, including biomass precursors (7).

Constraint-based modelling of genome-scale metabolic networks provides the means to *in silico* analyse microbial community interactions (8–12). The existing metabolic reconstruction approaches (13–19) rely on linking genome annotation to enzymatic reactions from various databases (19–23). A structural comparison of the metabolic models resulting from these approaches showed that the portions of shared reaction, metabolite, and gene sets are rather moderate (24). Hence, a consensus model could provide a means to combine the advantages of the existing approaches (25, 26).

However, a consensus model is not guaranteed to be functional (i.e. to simulate growth), as knowledge gaps may occur in all of the underlying (draft) models. To address this issue, several algorithms for gap filling, applicable to single models, have been proposed; however, they are not feasible for gap filling models of large communities due to the sheer amount of computational resources required (27, 28). Since members of microbial communities are often dependent on each other, gap-filling solutions of the community members must also be put into context of the community in which the organisms co-exist. In addition, the usage of gap-filled solutions and exudates from other community members may further reduce the overall number of added reactions to fill in the gaps in the metabolic model for the community. Existing computational efforts have shown that the gap-filling medium as well as the order in which gap-filling is applied plays a very important role in the reconstructed model (29).

Further, the problem of finding metabolic interactions within microbial communities using constraint-based methods has been addressed by multiple studies (8, 10, 30–34). However, most of these approaches are not suited for analysis of large microbial communities, comprising more than 10 microorganisms, due to the sheer size of the arising constraint-based problems. Importantly, the fully automated approaches rely solely on transport reactions in the model, and neglect permeability to define the set of exchangeable metabolites.

Here, we describe COMMIT, a constraint-based approach that respects the composition of the microbial community and metabolite leakage in the process of gap filling the consensus metabolic models of the respective community members. We then apply COMMIT to reconstruct high-quality metabolic models based for operational taxonomic units (OTUs) from the *Arabidopsis thaliana* microbial culture collections, *At*-SPHERE, for two different soil community compositions (35, 36). Altogether, our results demonstrate that the consensus approach in combination with the gap filling approach that respects the community composition renders COMMIT a valuable addition to the approaches for fully automated reconstruction of large-scale metabolic models of microbial communities.

## Results

### Reconstructed draft genome-scale metabolic models for 432 OTUs in *At*-SPHERE show substantial structural differences

We used the high-quality genomes for 432 OTUs available from *At*-SPHERE (37) to reconstruct draft genome-scale metabolic models. To this end, we applied four widely-used, fully automated approaches for metabolic network reconstruction relying on few parameters, namely: KBase (19), CarveMe (14), RAVEN 2.0 (17), and AuReMe/ Pathway Tools (15, 16) (for reviews, see (24, 38)). We then converted the draft models reconstructed by the four approaches to a common format by multiple technical adaptations (see Methods).

To compare the structure of the models after the conversion, we employed eight distance measures, including: the Jaccard distance based on the sets of metabolites, reactions, E.C. numbers, and dead-end metabolites, the number of dead-end metabolites, the rank correlation of E.C. number occurrence, usage of cofactors, the SVD distance of the stoichiometric matrices, and the Jaccard distance of gene identifiers (see Supplementary Methods). We then combined the resulting distance matrices in a compromise distance matrix for each OTU; to facilitate a global comparison of the different approaches, we also calculated the compromise distance matrix over all OTUs (see Methods, Figure 1A). The structural comparison revealed substantial differences across the draft genome-scale metabolic models generated by the four approaches (Figure 1A). In the compromise distance matrix obtained from the eight distance measures across all OTUs, the draft models showed an average distance of 0.64 to each other, ranging from 0.54 to 0.72 (with 1 denoting the largest difference). Regarding the gene identifiers, the models reconstructed using KBase, CarveMe, and RAVEN 2.0 were more similar to each other in comparison to the models resulting from AuReMe/Pathway Tools.

**Figure 1.**
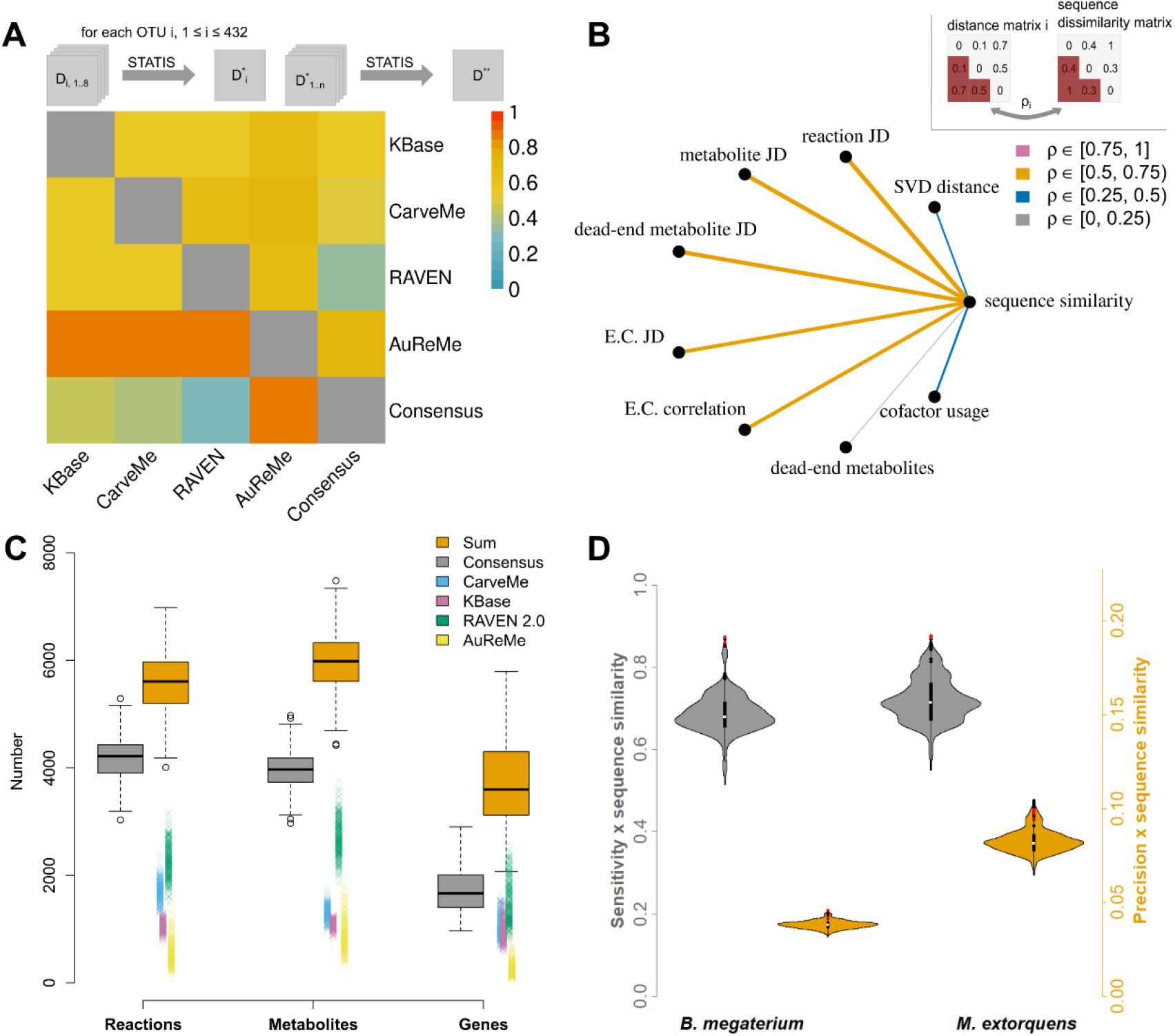
Structural comparisons of draft and consensus models. **(A)** Structural differences across the reconstructed draft models and the consensus. **upper triangle:** For each OTU, eight distance matrices (*D*) were computed to compare the models reconstructed using KBase (13), CarveMe (14), RAVEN 2.0 (17), AuReMe/ Pathway Tools (15, 16), and the consensus model. Detailed information on the distance measures can be found in the Methods section. These matrices were then combined per OTU by finding a compromise matrix 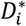 (STATIS method (59, 60)). The resulting matrices (one for each OTU, *n* = 432) were again combined using STATIS, resulting in an approach-by-approach distance matrix *D*^∗∗^. **lower triangle:** Gene Jaccard distances across all four methods and the consensus model. The compromise distance matrices were obtained as described above. Values close to zero indicate smaller differences. **(B)** Correlation of structural distances between the consensus models with sequence dissimilarity of 16S rRNA sequences. The same eight distance measures from (A) were used for pairwise comparisons of consensus models between all OTUs (*n* = 432). Detailed information on the distance measures can be found in the Methods section. Mantel correlation was used to compare each distance matrix to the sequence dissimilarity as depicted in the top-right corner. **(C)** Comparison of consensus models to draft models. The numbers of reactions, metabolites, and genes are shown for the consensus model in comparison to the single draft models generated by the four approaches. To depict the overlap between the four approaches, the sum of numbers of reactions, metabolites, and genes per model is shown (“Sum”, orange box). **(D)** Similarity of consensus models to selected reference models. The sensitivity (grey, left) and precision (orange, right) with respect to reaction, metabolite, E.C. number sets were calculated for each of the 432 models and each reference model. These values were scaled by the sequence similarity to the genome of the used references. OTUs that were assigned the same genus (9 for *B. Megaterium* and 27 for *M. extorquens*) or species (2 for *B. Megaterium* and 5 for *M. extorquens*) according to Bai et al. (37), are shown as black or red dots.

To determine whether or not the utilized distance measures are biologically relevant, we next calculated the correlation between the distance matrices and sequence dissimilarity of the 16S rRNA sequences. We found that the Jaccard distances corresponded to the generated phylogeny, with significant correlations ranging from 0.63 to 0.75 with an average of 0.70 (*p* < 0.001) over the 432 OTUs (Figure 1B). We also observed moderate but significant correlations of the SVD distance and the rank correlation of cofactor usage with sequence dissimilarity 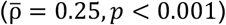. In contrast, the occurrence of dead-end metabolites showed a very low correlation with sequence dissimilarity 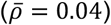. These findings indicated that the utilized distance measures to quantify the structural differences between the reconstructed draft models are biological relevant, further supporting our result that the models obtained by some of the utilized approaches are strikingly different (Figure 1A).

### Consensus metabolic models show high organism specificity

It has been demonstrated that the integration of multiple metabolic models into a consensus model leads to a reduced number of blocked reactions due to the complementarity of their information content (25, 39, 40). Since we observed that the draft models generated with the four reconstruction approaches differed in their underlying genome annotation and downstream reaction and metabolite sets (Figure 1A), we hypothesized that there would be an overall increase in model quality (assessed by the decrease in the number of gaps) of a corresponding consensus model. The consensus generation consisted of matching metabolite, reaction, and gene identifiers. Since the metabolites in the MetaNetX database are already structurally matched between various databases, duplicate metabolites could be removed from the consensus by only considering their identifiers. We employed cosine similarity to identify reactions of similar stoichiometry that may have opposite directions, lack protons or whose coefficients differ by a factor. Further, we compared mass-balance, reversibility, direction, and protonation. Previously published approaches as COMMGEN (40) or MetaMerge (25) were not applicable since they do not support the current MNXref format used in the MetaNetX database or are no longer maintained.

We found that the consensus model is considerably smaller than the mere sum for reactions, metabolites, and genes contained in the underlying draft models (Figure 1C). Further, the proportions of reactions, metabolites, and genes were not uniform across the draft models obtained from the different reconstruction approaches. The RAVEN 2.0 reconstructions consistently showed larger numbers of reactions, metabolites, and genes compared to all other methods, with AuReMe\Pathway Tools draft models exhibiting the lowest sums of these properties (Figure 1C). As a result, the draft models from RAVEN 2.0 exhibited the smallest overall distance (0.37) to the consensus model, while the draft models obtained from the other tools showed an average distance of 0.59 to the consensus (Figure 1A). The Jaccard distances of gene identifiers to the consensus model ranged between 0.31 for RAVEN 2.0 reconstructions and 0.87 for AuReMe models.

To show the organism-specificity of the resulting consensus model, we next assessed the similarity of the consensus to selected reference models for OTUs that were resolved down to species classification, namely *Bacillus megaterium* (iMZ1055) and *Methylobacterium extorquens* (iRP911). To this end, we assessed the manually-curated models for these two species (41, 42) by first translating their identifiers into the MNXref namespace (whenever possible), facilitating comparison of reactions, metabolites, and E.C. numbers. Every consensus model was compared to each of the two curated models to determine the number of true positives, false positives, and false negatives and to calculate sensitivity and precision (see Methods). Both values were scaled by the sequence similarity of an OTU’s 16S rRNA sequence to the sequence of the reference species. Despite the low translation efficiency, we found that the consensus models of OTUs of the same species were more similar to the curated models than models of OTUs that share only the genus or model of even less-related OTUs (Figure 1D). Notably, most of the models for OTUs with only matching genus exhibited higher precision for *M. extorquens* than the ones from the same species. Moreover, the consensus models showed higher values for scaled sensitivity compared to the individual draft models (Figure S1). In contrast, the values for the scaled precision of the consensus models were smaller than those of the draft models. This can be explained by the overlap of the draft models with respect to the sets of features (i.e. metabolites, reactions, and E.C. numbers): The number of the non-overlapping features (i.e. false positives) increases more than the number of features matching with the reference models (i.e. true positives) during the consensus generation. Hence, the precision is lower for the consensus as the ratio of true positives to false positives is shifted. Yet, across all draft models, except for AuReMe\Pathway Tools, models of the same species and genus were more similar to each other than models of phylogenetically more distant OTUs. Therefore, we concluded that the consensus of metabolic models shows a higher organism specificity as well as a higher quality than the draft models obtained from the individual approaches.

### COMMIT provides gap-filling that respect the community composition and metabolite leakage

Constraint-based analysis of microbial communities is based on assembling functional metabolic models for members of the community that can, in turn, be used to simulate microbial growth. However, obtaining a functional metabolic model from a draft model entails adding reactions to an incomplete metabolic network to enable simulation of growth (43, 44); often the inserted reactions are without genomic support.

The existing gap-filling methods are applied with models of individual species and do not consider the innate dependence between the community members (6, 9, 45). In contrast, our approach, termed COMMIT, aims to identify a minimal gap-filling solution that respects the composition of the microbial community. To this end, COMMIT explores a specified number of random orderings of the community members. Draft models for each member of the community are then gap-filled following a given random ordering of the community members. COMMIT relies on the FastGapFilling algorithm, which optimizes a weighted sum of the fluxes through the biomass and additional candidate reactions by using linear programming (LP) formulation (see Methods section for more detail).

More specifically, COMMIT starts with a minimal medium; exported metabolites from gap-filled models are then determined and used to enlarge the medium, thereby shaping the overall gap-filling solution. The random orderings are compared with respect to four criteria, including: (i) the number of added reactions, (ii) dependence of the first member on exported metabolites of subsequent models, (iii) number of exchanged metabolites, and (iv) the sum of biomass fluxes of all community members (see Methods). We determined exchangeable metabolites after gap-filling each of the draft models following the given random ordering of community members. A metabolite is considered exchangeable if its export does not decrease the flux through the biomass reaction by more than a pre-specified factor (e.g. 10%). We rely on using a threshold (of 10%) because an overly altruistic behaviour of single community members is not considered biologically relevant.

The set of exported metabolites is in turn made available to the subsequent models by allowing a lower cost for their import, thus enlarging the initial gap-filling medium (Figure 2A). Assuming that the community members depend on each other (6), we hypothesized that the enlarged medium is expected to reduce the added reaction sets of subsequent members of the community. Further, the existence of an optimal ordering is likely since gap-filling solutions do not only depend on the algorithm and candidate reactions employed, but also on the underlying medium (29). Given an ordering of microorganisms, the solution for the first model in the ordering is obtained by using its auxotrophic medium used in KBase (13). This strategy was employed to prevent an unrealistically high number of added reactions for the first model assuming the included compounds, most importantly essential amino acids, are likely produced by other community members. Altogether, the gap-filling procedure is conditional on the community composition, as it takes metabolic capabilities of all community members into consideration.

**Figure 2.**
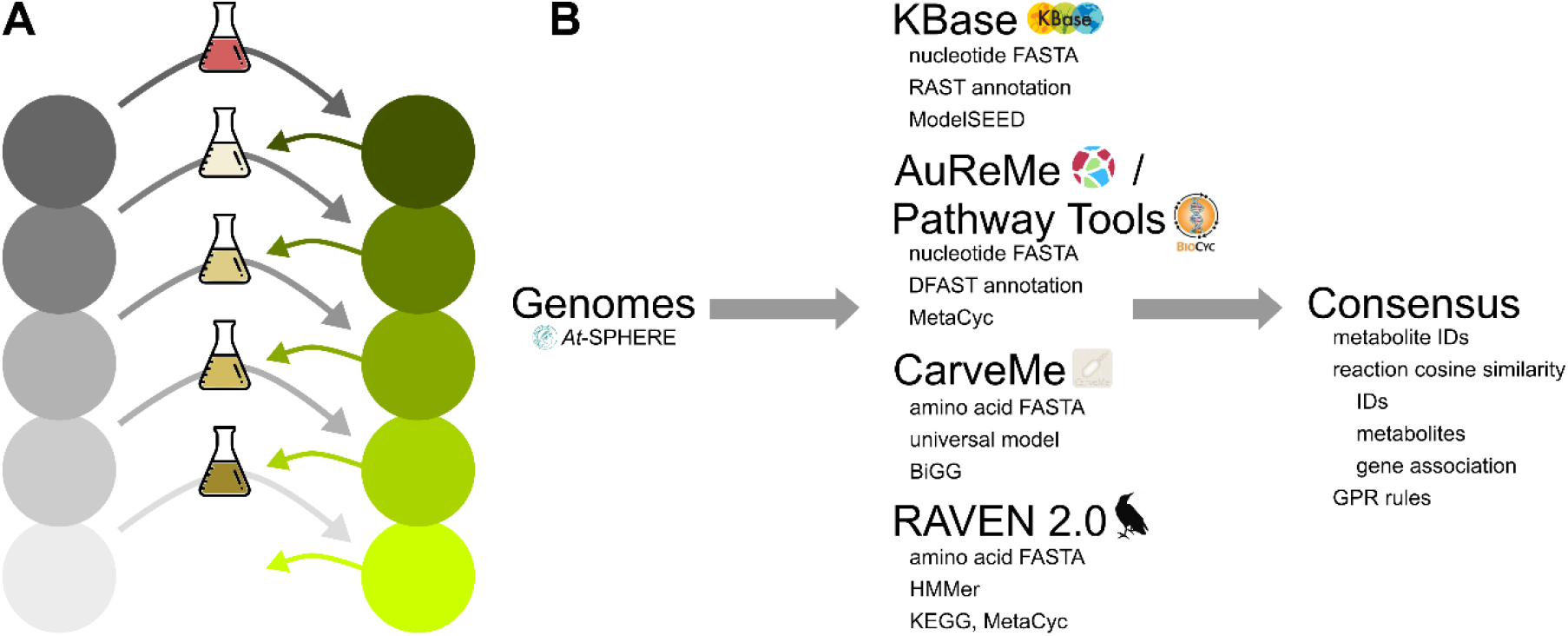
Schematic workflow of metabolic model reconstruction from multiple approaches using COMMIT. **(A)** The consensus models were gap-filled conditional on the community composition. Grey circles represent the models for every organism before gap filling. Green color indicates functional models after gap filling. The medium for the first model was the respective auxotrophic medium predicted using KBase (13) (red medium). For subsequent models, a minimal medium was used, which was enriched by the exported metabolites of already gap-filled models (green arrows). **(B)** Annotated genomes from *At-*SPHERE were used as the basis to reconstruct 432 draft metabolic models using the four recent methods from KBase (13), CarveMe (14), RAVEN 2.0 (17), and AuReMe/Pathway Tools (15, 16). These draft models were merged into consensus models per OTU.

In addition to the medium, the choice of candidate reactions affects the set of reactions that are added to close gaps in biological pathways (46). In COMMIT, the objective of the gap-filling LP includes specific costs for different reaction types (e.g. transport reactions for highly-permeable metabolites and reactions with sequence evidence). To predict metabolite permeability, we obtained molecular properties from PubChem (47) (see Methods section for details). Altogether, six parameters were used for permeability prediction based on Lipinski’s rule (48, 49), resulting in reduced costs for 2766 out of 4520 transport reactions in the gap-filling database. Moreover, reactions assigned to enzymes with sequence evidence in the respective genome were assigned a lower cost (see Methods).

### COMMIT reconstructs high-quality soil microbial communities based on consensus models from *At*-SPHERE

We applied COMMIT to two soil microbial communities whose composition was determined by comparison of 16S rRNA sequences from environmental samples with those from the cross-reference OTUs in *At*-SPHERE (37). This resulted in 20 and 24 recovered soil OTUs from two respective studies which, referred to as “Bulgarelli” (36) and “Schlaeppi” (35). The analysis of OTU abundances revealed the existence of high- and low-abundant OTUs in both experiments (Table S1, S2). The draft metabolic models for the members of the two communities were gap-filled by applying COMMIT with 100 iterations, corresponding to different random orderings, and an initial gap-filling medium containing only D-glucose as a carbon source (Table S3). The explored orderings were compared with respect to the number of added reactions, biomass fluxes, and number of exchanged metabolites, after scaling by their values in the optimal solutions (using the four abovementioned criteria).

Since solutions with the smallest number of added reactions across all iterations are generally favoured, all suboptimal solutions have a larger number of added reactions (Figure 3). Moreover, for both communities, we observed a shift in the number of exchanged metabolites when comparing the optimal with the suboptimal solutions from the other random orderings. While most of the suboptimal solutions exhibited a larger number of exchanged metabolites in the full consensus (Figure 3A,B), the peak shifts towards most solutions having a smaller number of exchanged metabolites in the consensus without CarveMe models (Figure 3C,D) and consensus only from KBase draft models (Figure 3E,F). This observation was likely due to the increasing importance of integrating more exchange reactions for metabolites provided by other community members to reduce the number of added reactions, when the number of merged models is lower. In contrast, such a pronounced trend could not be observed for the sums of predicted growth rates. Nevertheless, we observed that the distribution of scaled differences for the sum of biomass fluxes is similar to the distribution of exchanged metabolites for most of the simulations, suggesting a dependence between these two criteria.

**Figure 3.**
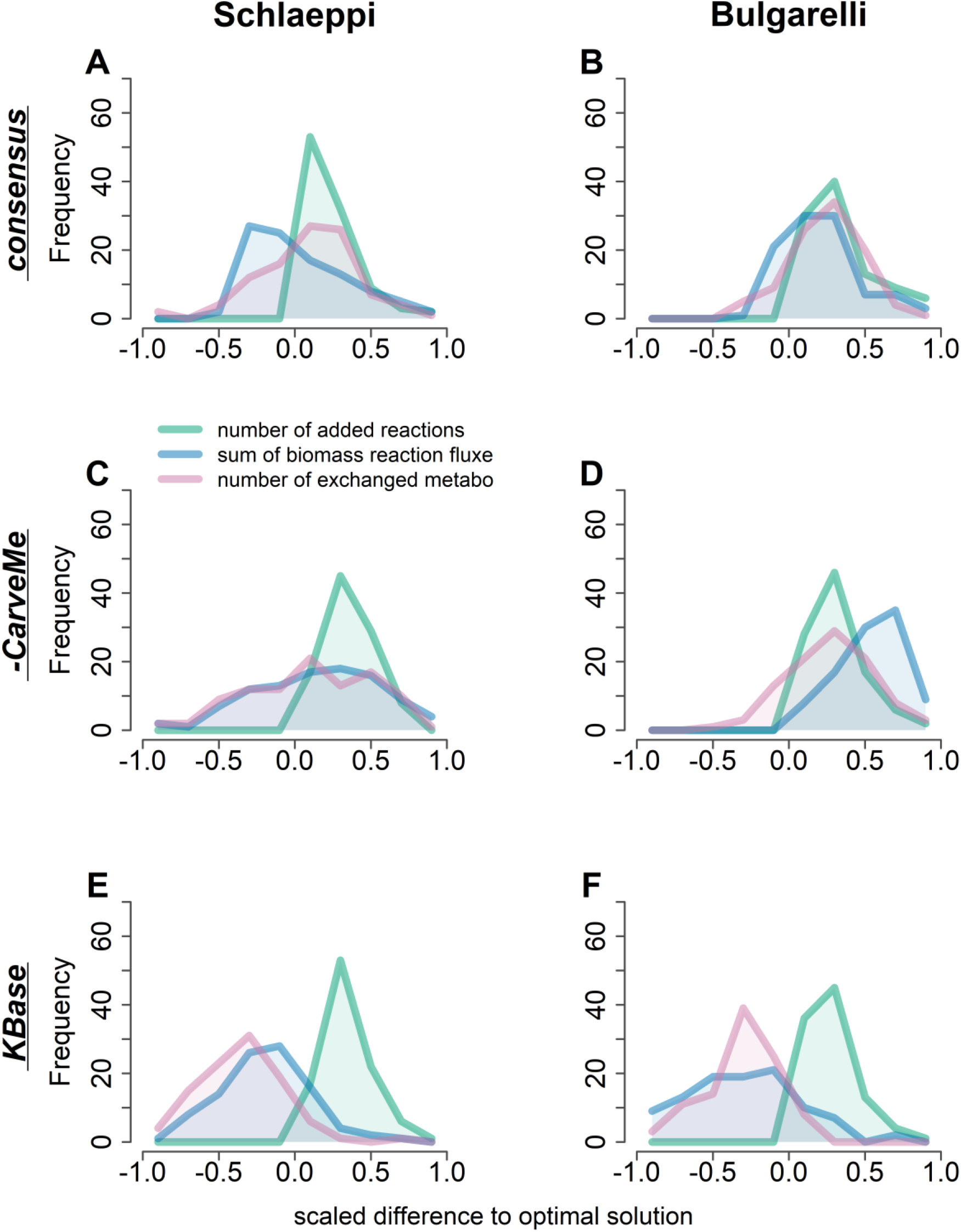
Differences across the random orderings of microorganisms explored in the gap filling that respect the microbial community composition. The number of added reactions, sum of biomass fluxes, and the number of exchanged metabolites were compared between all orderings explored by COMM**I**T. **(A**,**B)** consensus models from all approaches **(C**,**D)** consensus models without CarveMe draft models **(E**,**F)** converted KBase draft models. Results from the Schlaeppi community are shown on the left and results from the Bulgarelli community on the right panels. The lines represent the number of counts for each histogram bin, for the added reactions, growth rates, and numbers of exchanged metabolites. The values for each measure were scaled to the respective value associated with the optimal ordering. Number of iterations: *n* = 100; Number of OTUs per community: 24 for Schlaeppi and 20 for Bulgarelli.

We next used the *memote* test suite to assess the quality of the resulting functional models. The generated models of the Schlaeppi community exhibited an average score of 35% (Table S4). A strong reduction in the *memote* score was expected due to missing metabolite, reaction, and gene annotation by databases other than MetaNetX. Nevertheless, the generated models showed an average consistency of 44%. Importantly, the average fraction of reactions with associated GPR rules in the consensus was 90%, since reactions with genomic evidence were preferred during both the merging and gap-filling procedures. Therefore, the resulting models showed very similar characteristics to the previously used references models. In comparison, the reference model for *B. megaterium*, iMZ1055, was scored with 25% and 55% consistency.

### COMMIT shapes and significantly reduces the gap-filling solution

The observed differences between different random orderings of community members indicated that there exists an optimal solution that minimizes the number of added reactions for the particular community composition. Next, we were interested to assess the differences between the gap filling solutions for individual models with and without respecting the community composition. We found that COMMIT decreased the number of added reactions compared to the individual gap filling of the models using the same algorithm without allowing for metabolite exchange between the members (Figures 4A, S2). Despite the low number of added reactions for the consensus model, we observed a significant decrease in the gap-filling solution size with COMMIT in comparison to the classical gap filling approach (Figure 4A).

**Figure 4.**
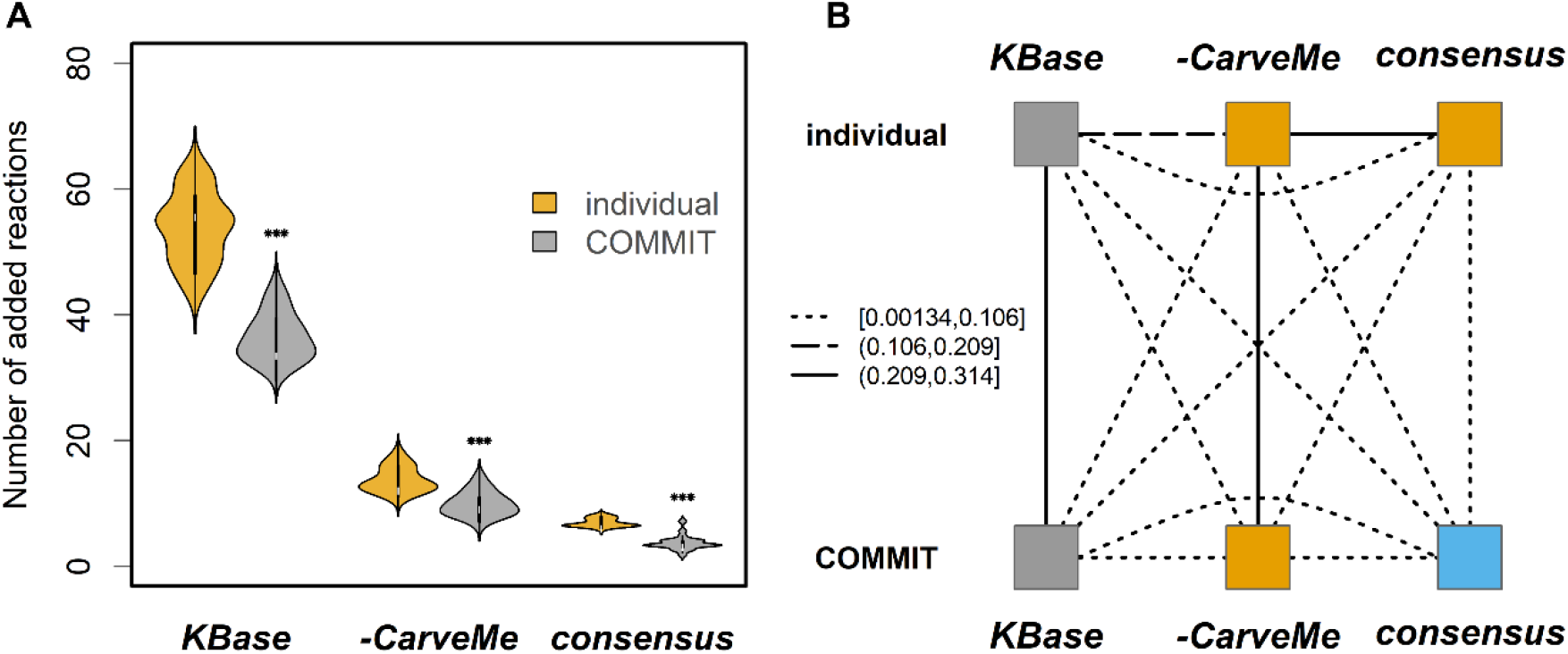
Comparison of gap filling solutions from individual and conditional gap-filling of consensus metabolic models in the Schlaeppi community. Full consensus (*consensus*), consensus without CarveMe models (*-CarveMe*), and KBase draft models (*KBase*) were gap-filled either individually or conditionally using the COMM**I**T approach. **(A)** Sizes of gap-filling solution sets were compared for each model type using a paired Wilcoxon rank sum test (∗∗∗ *p* < 0.001). **(B)** Pairwise comparison of added reactions obtained for each model and gap-filling type by calculating the Jaccard similarity per OTU. The resulting matrices were merged per group using STATIS method (59, 60). The obtained values were grouped using K-means clustering (*K* = 3). The line types indicate the average similarity between the compared groups.

To show that the set of added reactions is not only smaller but also differs in its composition, we performed K-means (*K* = 3) clustering with the matrix of Jaccard similarities of the added reaction sets for the full consensus, consensus without CarveMe, and KBase draft models gap-filled either with or without considering the community composition (Figures 4B, S3). We used three clusters based on the hypothesis that the solutions would group together by the model type (i.e. consensus, consensus without CarveMe, KBase draft models) rather than by the applied approach (i.e. with and without considering the community composition). We found that the reaction set added to the consensus models using COMMIT clearly separated from the other sets of added reactions, indicating that the result from the proposed approach differs substantially from those obtained by gap filling the models individually. The two other clusters were formed by KBase draft models and consensus without CarveMe models including the individual solution for the full consensus, respectively (Figure 4B).

### Exchanged metabolites reveal putative metabolic interactions

As the reduction of added reaction sets is based on the metabolic interactions between the models, the underlying set of exchanged metabolites was investigated to find putative dependencies within the investigated communities. To this end, we determined the contribution of each community member to the common pool of exchanged metabolites, without considering those that were already contained in the minimal medium.

First, we grouped the models based on bacterial families as found in *At*-SPHERE (37). As expected, members of all families show putative dependencies for carbohydrates, of which mostly β-D-galactose but also D-mannitol, maltotriose, maltopentaose, and β-D-fructofuranose were imported (Figure 5, Bulgarelli: Figure S4). In addition, amino acids, like: D-alanine, L-cysteine, L-leucine, L-asparagine, L-valine, L-isoleucine, L-arginine, and L-lysine were found to be exchanged. Further, we observed that members of the Micrococcaceae appear to have a rather broad spectrum of utilized nutrients, which spans most of the exchanged amino acids and carbohydrates. In contrast, bacterial families, such as: Xanthomonadaceae, Paenibacillaceae, and Mycobacteriaceae, were found to depend on a smaller, more specific set of imported metabolites. Although members of most of the bacterial families can export a variety of metabolites at a relatively low cost, there exist families that include more members exporting these metabolites (e.g. Mycobacteriaceae) than others (e.g. Micrococcaceae). Interestingly, cytidine, L-leucine, L-lysine, iminosuccinate, and *iso*-citrate were reported to be exchanged between only two families, respectively, indicating specialized interactions. In contrast, H^+^, P_i_, NH4^+^, Fe^2+^, Fe^3+^, and SO_4_^2-^ are exported at least from 20 out of 24 members. However, H^+^ import was observed for only 14 community members. This finding indicates that the members of a bacterial community prefer different pH in their local environment. This also suggests that almost all members have the ability to increase the acidity of the soil without a large growth compromise. Similarly, the import of P_i_, NH4^+^, Fe^2+^, Fe^3+^, and SO_4_^2-^ was observed in even fewer instances (*n* < 3).

**Figure 5.**
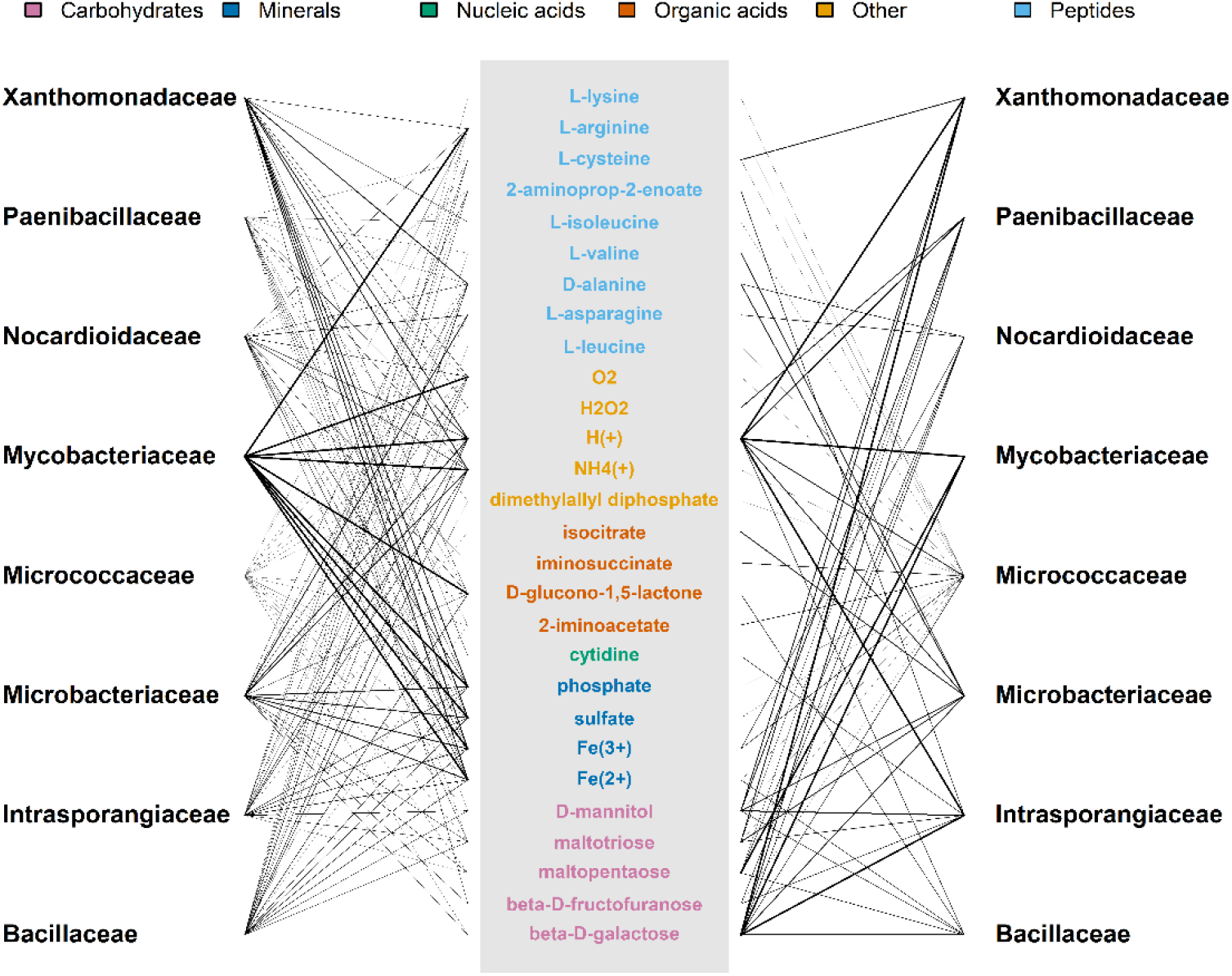
Putative metabolic interactions between the bacterial families. The sets of imported and exported metabolites that were determined for each member during conditional gap-filling and grouped into corresponding bacterial families. On the left-hand side, metabolite export is shown, which does not require a reduction of growth greater than 10%. The common pool of exchangeable metabolites is shown in the center, which were classified using the KEGG BRITE br08001 with manual refinement. On the right-hand side, import reactions are shown, which were introduced during the gap-filling procedure. Line widths represent the number of community members per import/export of a metabolite, scaled by the abundance of the respective family.

To analyse directed interactions between the community members in more detail, we conducted a downstream analysis of the gap-filled models using the SMETANA tool (10) (implementation from <u>github.com/cdanielmachado/smetana</u>). For the Schlaeppi community, we obtained a high metabolic interaction potential (MIP) of 0.98, which indicates that the community members can provide most of the essential nutritional components for itself by exchanging metabolites. SMETANA predicted metabolic interactions between 17 out of the 24 community members. To compare the obtained dependencies with our results, we transformed the directed interactions predicted by SMETANA to undirected interactions defined by the pairwise overlap of imported and exported metabolites. We found that about 21% of all possible interactions were predicted by both methods. Additionally, 22% of all OTU pairs were predicted to be non-interacting by both methods, summing up to 43% agreement between SMETANA and COMMIT (Figure S5). However, the intersection of the sets of exchanged metabolites from both methods consisted of only eight metabolites compared to a total of 28 metabolites from COMMIT and 22 from SMETANA; these included, aside from minerals, D-alanine, L-arginine, L-asparagine, L-cysteine, and NH4^+^.

To investigate the functional effects of the dependence on the import of metabolites, we investigated whether or not there is a decrease in growth upon blocking the respective uptake reactions for each metabolite. To this end, we calculated the ratio of biomass flux to the optimal growth rate without any constraint on exchange. In addition, we grouped the models based on these ratios using hierarchical clustering (Figure 6, Bulgarelli: Figure S6). The heatmap shows that resulting groups can partly be explained with the phylogeny, with some of the metabolic dependencies being independent of the taxonomic classification. Further, we observed that many OTUs strongly depend on β-D-galactose, maltotriose, and D-mannitol as carbon sources, while others prefer amino acids, such as: L-asparagine and D-alanine. Interestingly, two members of the genus *Paenibacillus* (i.e. Soil522 and Soil787) showed a strong reduction in growth when the import of protons or H_2_O_2_ is blocked. In addition, only one model assigned to the genus *Rhodanobacter* (Soil772) shows a strong reduction across both compositions when sulphur was not available. Indeed, it has been shown that a bacterium of the same genus, *Rhodanobacter thiooxydans* sp. nov. was found to potentially use sulphur as energy source for autotrophic denitrification (50). Therefore, our findings show that the fully automated metabolic reconstructions while considering the composition of microbial community and metabolite leakage results in high-confidence models, as corroborated by existing evidence.

**Figure 6.**
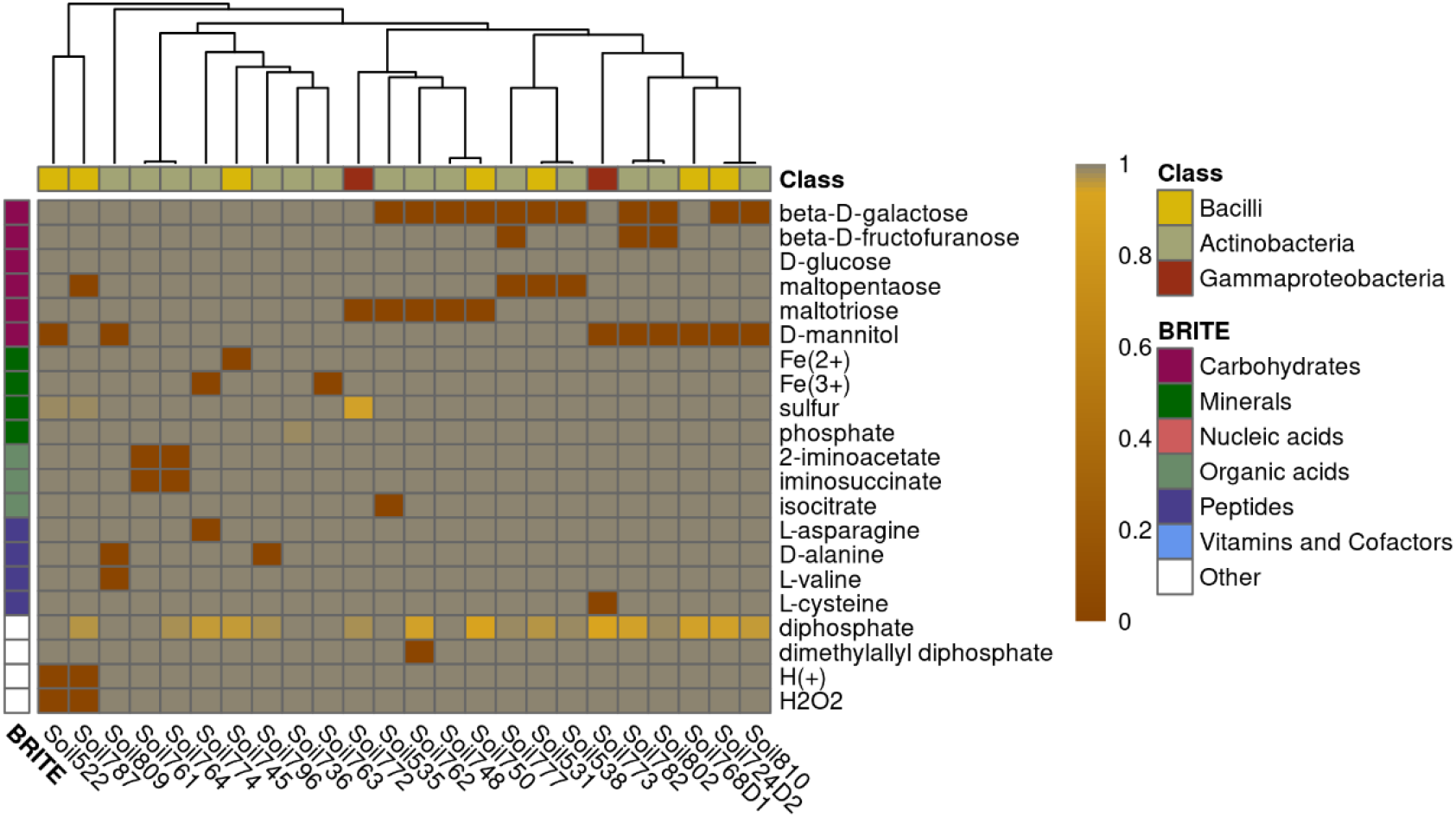
Reduction of biomass flux upon blocking the uptake of metabolites whose import was added during conditional gap filling. The uptake of every metabolite whose uptake was included during conditional gap filling was blocked separately and the growth rate under this condition was divided by the growth rate for that particular model. A threshold of 0.99 was applied to avoid false-positive responses. Biomass flux ratios were calculated for all metabolic models in the Schlaeppi community. The models were annotated to classes based on the taxonomy published by Bai et al. (37). The metabolites were grouped based on KEGG BRITE br08001, which was manually refined. Hierarchical clustering was performed using average linkage with Euclidean distance. Cells in dark red indicate low biomass whereas yellow cells indicate a smaller reduction of biomass.

## Discussion

Functional separation of the metabolic capabilities in a microbial community represents an important factor directing ecosystem functioning. While ecological interactions describe the effects that species can have on one another they miss out on complex metabolic relationships between multiple species. Leaky functions as metabolite diffusion within microbial communities can more likely explain the underlying metabolic interactions and dependencies (6). The investigation of communities is challenged by the number of participating members and the lack of detailed species-level information.

We investigated the usage of consensus metabolic models for assessing metabolic interactions between the members of a soil community. This was achieved by devising a novel approach, which fills gaps in the metabolic models while considering the community composition. More specifically, the obtained solution is dependent on the permeability-based metabolite exchange between the single models which allows for an investigation of their metabolic dependencies.

First, we demonstrated that the draft models generated using KBase (13), CarveMe (14), RAVEN 2.0 (17), and AuReMe\Pathway Tools (15) differ substantially from each other, as previously indicated by Mendoza et al. (24). We hypothesized that the consensus models generated from the draft models may resemble manually curated reference models of the same species. We showed that, despite exclusively using automated procedures, consensus models are highly organism-specific. Some of the models sharing only the same genus were structurally more similar to the reference than the ones which share the same species. Since in bacteria even strains within a species can differ from each other, the most similar models of the same genus could indeed also be of the same species or be similar due to metabolic niche adaptation. These results suggests that the draft models from different approaches complement each other. Further, pairwise Jaccard distances based on different model features and SVD distance based on the stoichiometric matrices correlated significantly with the phylogenetic distance. These results showed that the differences across the consensus metabolic models represent the phylogeny of the underlying OTU genomes, providing support for the biological relevance of the reconstructions.

The consensus models were then used as input to COMMIT to find a minimal set of reactions allowing growth simulations. As a result, COMMIT both shapes and significantly reduces the gap-filling solution. However, the set of added reactions by COMMIT was much smaller than that obtained by gap filling of individual models. In addition, we did not observe a significant correlation between the number of exported metabolites per model and the order in which models are considered for gap filling by COMMIT. Therefore, we concluded that the models that are gap-filled first, in the considered orderings of community members, provide specific compounds to the community that cannot be produced by other members. To corroborate this claim, we did not observe a reduction of biomass flux upon blocking of metabolites unique to the first five models in the optimal gap-filling order.

To confirm the putative dependencies, we investigated biomass reduction upon unavailability of each compound in the medium. Here, we observed that the metabolic dependencies of the models clustered in part according to phylogeny. Additionally, auxotrophies appeared to be not only influenced by lineage but also by specific community composition. This is in line with previous observations that auxotrophies can evolve relatively fast, given an environment or community providing the respective compounds, e.g. certain amino acids (51).

Finally, we assessed whether the given communities are, as hypothesized, divided into helpers and beneficiaries according to the Black Queen hypothesis (6). Even though the differences in exported metabolites were not large, it could be observed that members of the Mycobacteriaceae are among the OTUs that exported the most metabolites in both communities. Moreover, there were differences between the numbers of imported metabolites between bacterial families, which supports the existence of ‘helpers’ and ‘beneficiaries’ within the community.

By the community-dependent generation of functional metabolic models, we could show that prediction of metabolic capabilities as well as insight into the community structure can be obtained by solely using automated tools. In conclusion, this study shows that the usage of automatically-generated metabolic models provides a powerful means to analyse large-scale microbial communities even of uncharacterized species.

## Materials and Methods

### Genomic data

The 432 genomes that were used for genome-scale metabolic model reconstruction were downloaded from *At*-SPHERE (http://www.at-sphere.com) (37). The phylogeny information as well as the sequences from bacterial culture collections were also obtained from *At-*SPHERE. Abundances of the single OTUs in the environmental samples were computed using 16S rRNA sequences that were kindly provided by Ruben Garrido Oter (Max-Planck-Institute for plant breeding, Cologne). For this purpose, the USEARCH v11 software (52) was used to generate an OTU table, which was in turn normalized using the R implementation of the cumulative sum of squares (CSS) method (53). As an intermediate, the BIOM format (54) was used, and conversion was performed by the respective Python and R implementations.

### Draft model generation

The selected approaches, namely: CarveMe (14), KBase (13), RAVEN 2.0 (17), and AuReMe/ Pathway Tools (15, 16), use different annotation methods and databases which increases the variability of the automatically reconstructed models. For the CarveMe and RAVEN 2.0 draft reconstructions, the structural annotation from *At*-SPHERE was employed. CarveMe uses a manually curated, universal bacterial model in which BiGG reactions are linked to sequences. Upon sequence comparisons to the provided reference, reactions with low sequence similarity score of their associated enzymes are removed from the universal model. This approach, thus, guarantees an initial functional model given the medium that was used during the reconstruction. Moreover, draft models reconstructed with CarveMe contain a universal biomass reaction, as suggested by Xavier et al. (55).

The RAVEN 2.0 reconstruction pipeline for KEGG-based models constructs multiple sequence alignments (MSA) for all KEGG Orthology (KO) sequences, which were filtered for phylogenetically close sequences. Hidden Markov Models (HMM) were computed based on the MSAs and subsequently searched for in the genome using HMMer v3.2.1 (56). In addition, a MetaCyc-based model was generated based on sequence comparison to a complete model of the MetaCyc reaction database. Both model types were created and merged for every OTU using functions of the RAVEN 2.0 toolbox.

To reconstruct draft models using KBase, the genome sequences were re-annotated using RAST (57) with default parameters. To this end, we used the nucleotide sequences of the assemblies. The functional annotation from RAST is linked to reactions in the ModelSEED reaction database, allowing for the reconstruction of a draft metabolic model. The draft models were then downloaded from KBase for further processing.

Further, AuReMe/Pathway Tools was used as a fourth reconstruction method. These models were generated based on data files generated by Pathway Tools (16). As a functional annotation is required as an input for the Pathway Tools pipeline, all genomes were annotated using DFAST (58) using default parameters. The resulting draft models from AuReMe contained reactions based on the MetaCyc database.

### Model comparison

Metabolic models can be compared structurally using similarity measures that either use the stoichiometric matrix itself of particular features of the model (e.g. reaction or metabolite identifiers). The definitions of all distance measures that were used are described in the Supplementary Methods. First, the distance matrices were combined per OTU by finding a compromise matrix using the STATIS method (59, 60). Additionally, the Jaccard distance of gene identifiers was calculated between all four methods and the consensus for each OTU. As a second step, all resulting gene Jaccard distances and compromise matrices were again joined using STATIS (59, 60).

### Generation of consensus models

As a first step, the draft model from all approaches were converted to have the same number of fields, which in turn share a common format. This is particularly important for reactions, metabolites, genes, and gene-protein-reaction (GPR) rules. Reaction and metabolite identifiers were translated to MNXref identifiers using the MNXref reference files for metabolites and reactions. If a periplasmic compartment was present, all metabolites and reactions were treated as extracellular. Gene identifiers of draft models reconstructed using KBase and AuReMe/ Pathway Tools were translated by BLAST+ (61) search against the respective structural annotation obtained from *At*-SPHERE. For unification reasons, also biomass and exchange reactions, if present, were removed.

### Comparison to reference models

The cross-reference OTUs in *At*-SPHERE (37) contained isolates with species resolution, among them *Bacillus megaterium* and *Methylobacterium extorquens*, for which manually-curated models had been published. The models for *B. megaterium* WSH-002 (42) and *M. extorquens* AM1 (41) were downloaded and their features were translated to MNXref name space whenever possible. The comparison was hampered by the lack of cross-references to name spaces other than self-defined identifiers. True positives were defined as those identifiers of the consensus model that were also present in the respective reference. In contrast, false positives were defined as the difference between the consensus and the reference model. Conversely, the difference between the reference model and the consensus comprised the false negatives. Since the true negatives could not be determined, only sensitivity and precision were calculated as measures to assess the quality of model prediction.

The genomes of the two reference models were downloaded from NCBI reference numbers <u>CP003017</u> and <u>CP001510.1</u>. MAFFT online service (62) was used to generate MSAs of the 16S rRNA sequences of the OTUs and the respective reference, and for NJ/UPGMA phylogeny computation to generate a Newick tree that was converted into a distance matrix using Newick Utilities (63). Both MAFFT and Newick Utilities were used with the default parameter set.

### Biomass reaction and media

After the merging draft reconstructions from the four different approaches, a universal biomass reaction was added to the models. The metabolites and their coefficients in the biomass reaction were adopted from the CarveMe reconstructions (14) as they include the universal biomass reaction suggested by (55). In addition, the initial medium was considered by adding exchange reactions to the models. The initial medium originates from the intersection of predicted auxotrophic media for the given community obtained from KBase, without organic compounds (except D-glucose as single carbon source).

### Prediction of permeability

To predict metabolite permeability, chemical properties, namely:molecular weight (MW), polar surface area (TPSA), number H-bond donors (HBD), number of H-bond acceptors (HBA), rotatable bonds (RB) and predicted octanol/water partition coefficient (XLogP3), were obtained from PubChem (47) for all metabolites in the MNXref database (Figure S7, S8). This was achieved by translating a unique set of 451219 associated InChI keys to PubChem CIDs, whenever possible (116458 out of 451219: 26%), so molecular properties for 110266 compounds could be obtained. Since not all entries had an associated XLogP3 value (only 11894: 11%), the missing values were predicted using k-Nearest-Neighbor (kNN) regression with *k* = 1 using MW, TPSA, HBD and RB as predictors (Figure S9) (10-fold cross validation (CV): 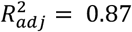); *RMSE* = 126) (validation set: 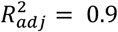; *RMSE* = 1.11). The value for *k* was determined by comparing the performance of multiple kNN models with 1 ≤ *k* ≤ 20 (Figure S9).

The chemical properties were used to classify metabolites into likely permeable and otherwise by applying Lipinski’s rule of five and including RB and TPSA (48, 49). Precisely, the applied rules were: HBD ≤ 5,HBA ≤ 10, MW ≤ 500 Da, TPSA ≤ 140 Å, *RB* ≤ 10, −0.4 ≤ XLogP3 ≤ 5.6.

If no properties were available, the respective metabolite was classified as not likely to be permeable. The above-described classifier was applied to all metabolites in the gap-filling database, which is depicted in more detail below. In total, 8354 out of 33547 (24.9 %) metabolites were predicted to be likely permeable. This set of highly-permeable metabolites was enriched with carbohydrates and fatty acids, but also nucleic acids and peptides (*p* < 0.001, ChEBI metabolite ontology (64)). The p-values were corrected for multiple testing using the Benjamini-Hochberg procedure (Table S5). Out of 4520 transport reactions, 2766 included at least one highly-permeable metabolite.

### COMMIT formulation

To perform gap filling of the generated models we implemented he FastGapFilling algorithm (65) in MATLAB and used specific weights for transport reactions (100), metabolic reactions (non-transport, 50), changing a reaction’s directionality (25), reactions including highly-permeable metabolites (subset of transport reactions, 50), reactions with sequence support (25), allowed uptake reactions for metabolites that are exported by other members (1) and exchange reactions (10^5^). The penalties applied in this study resulted from the comparison of different weights that were altered until satisfying results were obtained for selected models. The upper limit for the biomass flux was constrained to 2.81 *h*^−1^ as this is the observed growth rate for very fast growing bacteria as *Vibrio natriegens* (66). The latter was used to avoid the flux through the biomass reaction being pushed to its default limit of 1000, which would allow a higher number of reactions to be added. The additional constraint on biomass is, however, not propagated to the resulting model, as it is only part of the gap-filling LP.

The chemically-balanced part of the MNXref database was used as a gap filling database, including 25,740 reactions and 33,547 metabolites, after removing reactions according to the criteria described below. To this end, all generic compartments were ignored since a unique translation to either cytosol or extracellular space was not possible. Further, the metabolite ID ‘MNXM01’ was replaced with ‘MNXM1’ since both encode *H*^+^. Additionally, export reactions i.e. reactions that include metabolites associated to the ‘@BOUNDARY’ compartment were excluded from the gap-filling database. Reactions including stoichiometric coefficients above 20 were also removed.

To determine weights that quantify sequence similarity, HMMs constructed for all KO files were queried against every genome using HMMer (56). The KO identifiers were matched to MNXref reactions via associated KEGG reactions. An E-value of 10^−6^ was used as a cut-off. A lower weight was also assigned to transport reactions that include highly-permeable metabolites as indicated above. To this end, a subset of transport reactions that use highly-permeable metabolites was found in the gap-filling database. Further, one can choose to include sink reactions for cytosolic metabolites in the objective of the gap-filling LP. An additional matrix for sink reactions *R*_*sink*_ is then created for metabolites that are predicted to be highly-permeable or take part in a transport reaction in the model. These sink reactions can also be assigned a specific weight. Since a weighted sum of biomass flux and flux through the added reactions is maximized, only those sink reactions will be added that do not largely decrease the biomass flux. The sink reactions that were found during gap-filling will not be added to the model. Instead, export reactions from the cytosol to extracellular space as well as exchange reactions for the respective extracellular compounds are added to the model.

For the conditional gap filling, 100 random orderings of the considered models were inspected during COMMIT. The goal is to identify the best ordering with respect to specified criteria. More specifically, the models are gap-filled following a given ordering. To this end, every iteration starts with a minimal medium, which is augmented by the exportable metabolites of the gap-filled models. The medium for the first model in each iteration comprises metabolites that render the particular model auxotrophic. This is used so as to avoid the gap filling of the first model in the ordering to be unrealistically large in comparison to solutions for subsequent models. The auxotrophy media were obtained from KBase using the corresponding function. To arrive at a minimal medium, only nutrients were considered that were found in all auxotrophic media of the considered organisms, excluding amino acids and carbon sources, except for glucose. After each single gap-filling run, exportable metabolites are predicted and made available to the subsequent models via additional uptake reactions in the gap-filling database. The prediction of exchanged metabolites can be done during the gap filling step of the individual model or right after. When the first option is used, sink reactions for cytosolic metabolites are integrated directly into the objective of the gap-filling algorithm. Otherwise, a similar LP is solved using the gap-filled model maximizing the flux through sink reactions without reducing the biomass production by more than the given factor. For the second option, the optimal biomass flux 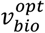 is determined first, followed by the maximization of flux through the sink reactions while guaranteeing a sub-optimal biomass 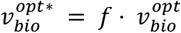. In this study, *f* = 0.*9* has been used (please see supplementary Methods for more detail).

Linear Program for the inclusion of sink reactions after gap filling:

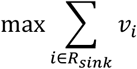

**s.t**

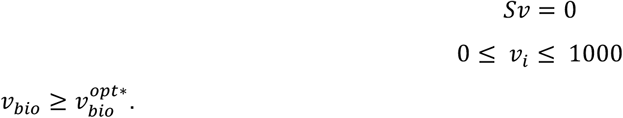

Please, refer to the Supplementary Methods for the full specification of the COMMIT procedure.

## Acknowledgments

P.W. and Z.N. would like to thank the Research Focus Group “Evolutionary Systems Biology” of University of Potsdam for funding. Z.N. would like to thank the Max Planck Society for funding.

## Author contributions

Z.N. and P.W. designed the research. P.W. performed all simulations and data analyses. Z.N. and P.W. wrote the manuscript. All authors read and approved the final manuscript.

